# NJGPT: A Large Language Model-Driven, User-Friendly Solution for Phylogenetic Tree Construction

**DOI:** 10.1101/2024.12.02.626464

**Authors:** Zhixuan Wang, Haoyuan Huang, Teng Li, Allen Rodrigo

## Abstract

**Motivation:** Phylogenetic reconstruction plays an integral part in much of the research in evolutionary biology. Currently, a newly minted phylogeneticist must choose amongst a relatively large array of phylogenetic software, each often with its own analytical routines, inputs and outputs. Our overarching aim is to construct a user-friendly pipeline with the most recent generative AI tool, ChatGPT-4 (at the time of writing) released by OpenAI, that is able to understand queries written in natural language, to build a phylogenetic tree using sequence data. By doing this, we demonstrate how generative AI may be used in phylogenetics, as a proof-of-concept. We also demonstrate the steps needed, presently, to ensure that ChatGPT can build phylogenetic trees accurately.

**Results:** We present NJGPT, a phylogenetic tool built using ChatGPT, a Large Language Model (LLM) Generative Pre-trained Transformer (GPT),which employs the Neighbor-Joining method to construct phylogenetic trees. NJGPT simplifies phylogenetic tree construction by allowing users to generate and visualize trees using natural language queries. It supports multiple sequence file formats, matrix calculation models, and gap-deletion methods. To evaluate the performance of NJGPT, we compared output and runtimes with the widely-used phylogenetic software, MEGA. Our results show that NJGPT produces identical trees over a range of sequence lengths and simple models of evolution. However, NJGPT faces visualization issues with datasets over 50 taxa and operational failures with larger datasets due to token limits. NJGPT runtimes were also substantially slower than MEGA; however, NJGPT’s user-friendly interface makes it ideal for beginners.

**Availability:** This plugin is available for free at https://chatgpt.com/g/g-1OzP3Qviw-njgpt, the source code is available on GitHub (https://github.com/ZWan622/NJGPT1.0.git) and is implemented using Python

**Supplementary information:** Supplementary data are available at *Bioinformatics* online.

## 1 Introduction

Phylogenetics is an essential branch of biology that models and analyzes the evolutionary relationships and histories of species, genes, languages, cultures, and other groups exhibiting variation that may be accounted for by shared ancestry and independent evolutionary change (Munjal et al., 2018). Phylogenetic reconstruction results in the construction of a bifurcating or multifurcating tree, which may be rooted or unrooted, with the terminal nodes of the tree representing the evolutionary units under study, and the internal nodes representing hypothetical ancestors (Randall et al., 2016; Wake, 1978).

Numerous phylogenetic tree construction programs have been created using various algorithms and data types to accommodate the increasing complexity of phylogenetic methodologies. These include the Neighbor-Joining (NJ) method, a distance-based clustering algorithm (Saitou & Nei, 1987); the Maximum Likelihood (ML) method, a probability-based algorithm that estimates the tree(s) most likely to give the observed data under a model of evolution (Felsenstein, 1981); parsimony algorithms that minimize the number of changes on a tree required to account for the observed data (Fitch, 1971); and Bayesian inference, which delivers probability distributions of trees and attendant parameters (Rannala & Yang, 1996). Biological data used in phylogenetic analyses include nucleotide sequences (Lewis, 1998), morphological data (Brazeau et al., 2018), protein-coding genes (Fong & Fujita, 2011), biallelic markers such as SNPs and AFLPs (Zhu et al., 2017), and many other datatypes.

Examples of phylogenetic software include PAUP*, which implements parsimony, ML, and distance-based methods including NJ (Wilgenbusch & Swofford, 2003); MEGA, which constructs trees using NJ and ML algorithms (Newman et al., 2016); and BEAST, which employs Bayesian algorithms to reconstruct posterior probability distributions of trees and attendant model parameters (Bouckaert et al., 2014). Although these programs typically have user-friendly interfaces, they require users to be familiar with the features of the programs, specific terminology, and the order of each processing step, and this maybe cumbersome for beginners to navigate, thereby significantly raising the barrier to entry. Additionally, some software can only read specific file formats, complicating data processing. Overall, existing software may not be as user-friendly as one would like, for researchers new to phylogenetics.

Artificial Intelligence (AI) with natural language capabilities that use generative pre-trained transformers (GPTs), like ChatGPT (www.openai.com), Gemini (www.gemini.google.com), and Copilot (www.copilot.microsoft.com), are being used increasingly in scientific research (Bano et al., 2023). Natural language GPTs typically use highly parameterized Large Language Models (LLMs) to model contextual relationships and co-occurrence amongst semantically meaningful fragments of text (these are called “tokens”, and may represent whole words, parts of words, e.g., suffixes, prefixes, characters and punctuation marks). Recently developed GPTs excel in generating and summarizing text (Cai et al., 2023); for instance, arguably the most popular GPT, ChatGPT, has a friendly interface and conversational capabilities that provide an accessible entry point for users new to AI and complex technical tasks (Sakirin & Ben Said, 2022). Notably, more sophisticated, recent highly parameterized models of ChatGPT-4 excel at coding, reviewing, working on images, and debugging computational code (Rahman & Wong, 2023). GPT-4’s code interpreter feature allows users to run simple code directly within its interface (Low & Kalender, 2023). Moreover, customizable GPT, a recent innovation by OpenAI, allows developers to create AI assistants by using natural language prompts to develop tailored versions of ChatGPT (OpenAI, 2023). Developers can build, share, and use these assistants for specific needs, integrating external data and Application Programming Interface (API) calls for personalized tasks, with all dialogue privacy protected.

However, controversy remains about ChatGPT’s ability to perform viable phylogenetic analyses. Shue et al. (2023) found that while ChatGPT-3.5 can generate workable code for unrooted trees using multiple alignments of protein-coding sequences of the TP53 tumor suppressor gene, it produces irrelevant functions for more complex tasks when using a designated species as an outgroup to root the tree, requiring manual intervention. To address these gaps, we created NJGPT by implementing the NJ (Neighbor-joining) algorithm (Saitou & Nei, 1987) in a customized GPT to make tree-building software more novice-friendly. NJGPT, implemented in ChatGPT, is the first phylogenetic tree-building software to run on a GPT platform. It responds to input files and user queries in natural language, aiming to lower the barrier for phylogenetic tree reconstruction, regardless of users’ programming and phylogenetic backgrounds. We have developed NJGPT as a proof-of-concept to illustrate how developers can leverage ChatGPT’s natural language processing capabilities to make phylogenetic tree construction more intuitive and accessible, simplifying the user interface and processing steps, and visualizing tree topology within the ChatGPT dialog interface.

## 2 Tool Description

The NJGPT pipeline is shown in Figure 1A. With ChatGPT’s assistance, we developed 11 Python-edited phylogenetic tree computation code files and integrated them into NJGPT using the GPT creation function. The names of the block files and the associated functions are shown in Supplementary Table 1. NJGPT runs these code files in a sandbox environment, processing and analyzing user-uploaded data based on our instructions (Low & Kalender, 2023). NJGPT uses ChatGPT’s programming capabilities to generate code for reading user-uploaded alignment files (e.g., FASTA, PHYLIP, MEGA, and XML) and extracting and displaying sequences. NJGPT presents six options: two deletion methods (complete and pairwise) and three models of evolutionary distance: Hamming distance, the Jukes-Cantor (JC) model, and Kimura two-parameter (K2P) model. Users can ask NJGPT about these models, including definitions and differences of pairwise and complete deletion. Each choice of model triggers the corresponding six code blocks in the NJGPT backend, based on the user selections. The remaining code blocks are then executed to compute the final phylogenetic tree. Users receive the corresponding tree files in Newick format and phylogenetic tree pictures with taxon names and branch lengths.

**Figure 1.**
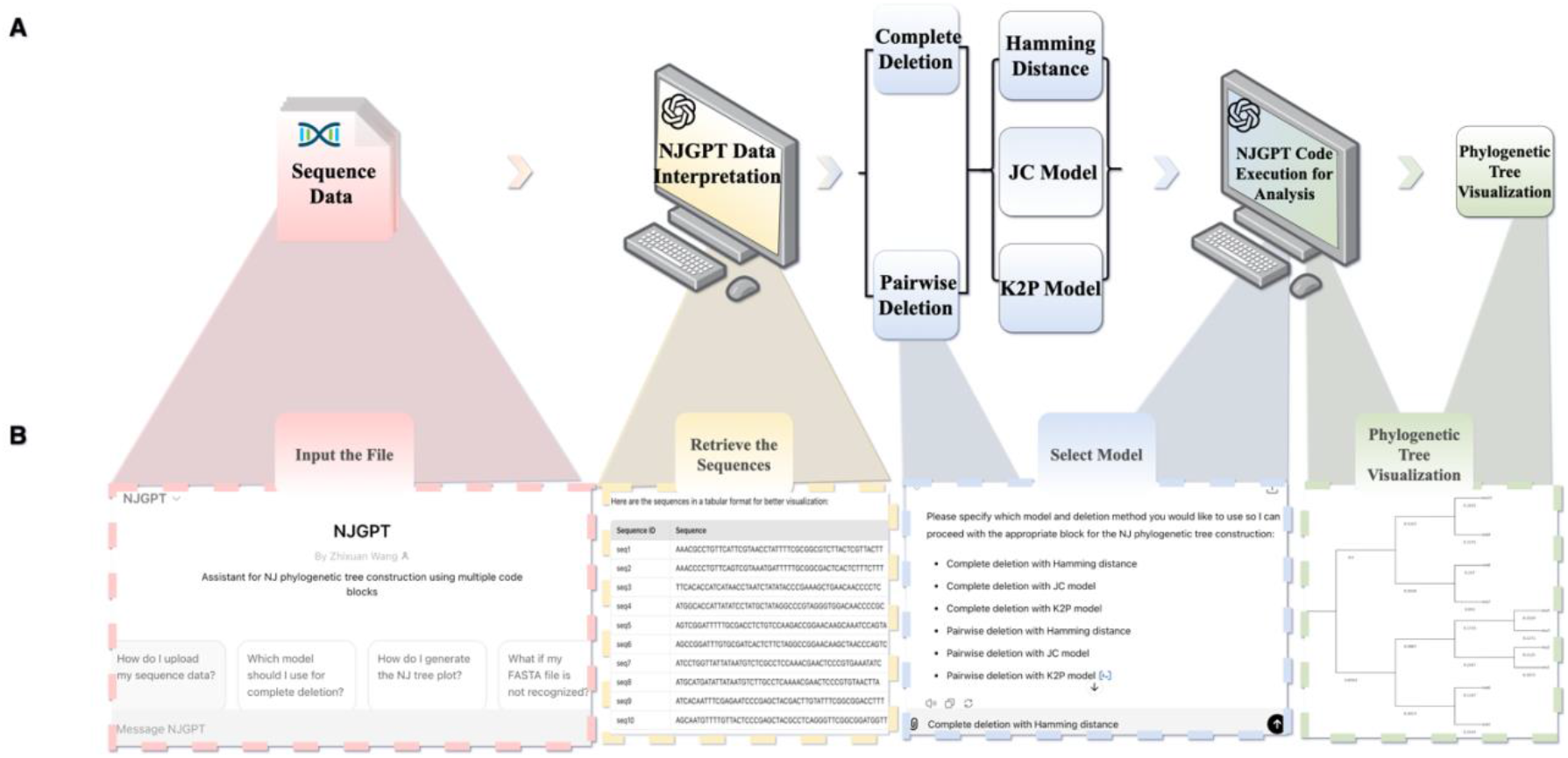
Preview of the pipeline and workflow of NJGPT. (A) The pipeline of NJGPT uses the code related to the NJ algorithm to run on ChatGPT. (B) The simple and user-friendly workflow of NJGPT, with corresponding outputs.

Figure 1B shows the user workflow. Users can open NJGPT in ChatGPT, access the interactive interface, and input the file they wish to analyze in the chat box. NJGPT reads the corresponding file and then prompts the user to choose a model. After the model selection, NJGPT runs code to calculate and visualize the final phylogenetic tree. Users can ask NJGPT about phylogenetic terminology and usage questions, and it will provide explanations in a user-friendly manner. If data cannot be processed, NJGPT will provide a pipeline with instructions on how to handle it.

## 3 Assessing Accuracy and Performance of NJGPT

The computational accuracy of NJGPT was evaluated using simulated data. Seq-gen (Rambaut & Grass, 1997) was utilized to generate a set of test alignments of evolutionarily related sequences using the HKY model, with lengths of 50, 100, 500, and 1000 nucleotides, and varying numbers of sequences of 10, 30, and 50. All possible combinations of these lengths and quantities were tested. The assessment focused on comparing the tree topologies and branch lengths generated by NJGPT with those from MEGA v11.0.13 based on the NJ method, employing various gap-deletion methods and distance models (Stecher et al., 2020; Tamura et al., 2021). The *phangorn* package in R (Schliep, 2011) was employed to calculate the Robinson-Foulds (RF) distance and patristic distance between the trees, phylogenetic trees, especially with increased sequence lengths. We believe that this is offset by an intuitive and user-friendly interface that may better suit users who value the ability to ask questions or guide a particular analysis using simple language. Consequently, although MEGA with these distances used respectively to compare the topological similarity and branch lengths of the phylogenetic trees generated by NJGPT and those from MEGA. For all simulation datasets, the time NJGPT required to construct phylogenetic trees was recorded and compared with the performance of MEGA.

## 4 Results

Across all simulations, the RF distances and patristic distances between the NJGPT trees and the MEGA trees were zero. Thus, in all tested scenarios, involving various deletion methods and distance models, the phylogenetic trees produced by NJGPT were identical to those from MEGA in terms of topology and branch lengths. This is illustrated using the MEGA example file “hsp20.fasta” in Supplementary Figure 1.

NJGPT reliably generates phylogenetic trees for datasets with up to 50 taxa and 10000 bases, producing clear images without any graphical artifacts. However, when the number of taxa exceeds 50, visualization issues emerged, characterized by overlapping taxa and branches, despite the production of accurate Newick format tree files. Performance degraded with larger datasets: the generation of Newick files became sluggish at 200 sequences, and NJGPT encountered operational failures for datasets with 400 taxa and sequence lengths of 5000, due to exceeding token limits in the web version of ChatGPT (OpenAI, 2024). These limitations manifest as “execution environment” errors. Tokens pose constraints on the tool’s ability to manage large datasets, significantly impacting its performance (Liu et al., 2023; Sun et al., 2023).

Runtime comparisons between NJGPT and MEGA (Supplementary Fig. 2) revealed that NJGPT generally requires more time to generate may be preferable for those requiring faster computations, NJGPT is distinguished as a more accessible tool for beginners.

## 5 Conclusion

As a proof-of-concept, we have used a relatively simple phylogenetic tree reconstruction algorithm to demonstrate that LLM GPTs can be used to develop tools appropriate for phylogeneticists. As the first tool to compute phylogenetic trees within ChatGPT’s web interface, NJGPT integrates ChatGPT’s natural language processing capabilities with the Neighbor-Joining (NJ) algorithm to provide an easy-to-use tool for constructing phylogenetic trees. NJGPT can process most used phylogenetic file formats and interactively guide users through the process, offering explanations and recommendations.

Despite NJGPT’s strong performance with most simulated datasets, it has some drawbacks: computation time is relatively long, usually ranging from one to three minutes, and increases with sequence complexity. When the number of sequences reaches 50, branches on the phylogenetic tree image may overlap, and with more sequences, the overlap becomes more severe. When the number of sequences exceeds 200, NJGPT cannot perform calculations because of the large data volume. Additionally, NJGPT cannot perform sequence alignment independently, requiring an alignment file as input.

Nonetheless, we believe that NJGPT lowers the entry barrier to phylogenetics, making it easy for researchers to generate and visualize phylogenetic trees regardless of their programming and phylogenetic background. This innovation facilitates the application of phylogenetics in various research fields and opens new possibilities for AI’s widespread use in scientific research. It may also be useful as a teaching aid for phylogenetic studies.

At present, these tools like NJGPT need to be assembled using a mixture of code, potentially developed with the assistance of ChatGPT and parsed through ChatGPT’s code interpreter, complemented by ChatGPT’s natural language capabilities. We anticipate that as LLM GPTs improve, there will be less and less need to provide code to execute some of these phylogenetic algorithms, however, the extent to which LLM GPTs will be able to deal with the more mathematically complex phylogenetic solutions remains unknown and should be tested repeatedly as newer versions of LLM GPTs emerge.

## Acknowledgements

We extend our gratitude to the ChatGPT and OpenAI team for their contributions to the GPT functions, and to Andrea Grecu, June Ko, Miao Wang, and Yuan Xu, for their invaluable support and discussion. Beyond using ChatGPT-4 to assist with the development of NJGPT, ChatGPT-4 was also used in the drafting of this manuscript to provide language translation and editorial assistance, including corrections of grammar and syntax. Research design, and conceptual content are solely the responsibility of the authors.

