## Supplementary Table1, Table2, Figure1 and Figure2 for "NJGPT: A Large Language Model-Driven, User-Friendly Solution for Phylogenetic Tree Construction"

**Supplementary Table1.** Summary of the functions of various scripts used in the process of constructing phylogenetic trees based on the Neighbor-Joining method. Each script is responsible for specific steps and methods, ranging from calculating sequence distances to generating and visualizing the final phylogenetic tree. Distance calculations are conducted using either complete deletion or pairwise deletion methods, accompanied by various models such as the Hamming Distance, Jukes-Cantor (JC) model, or Kimura 2-parameter (K2P) model. Subsequent steps (Block2 through Block6) entail processing and analyzing the sequence data to construct and visualize the Neighbor-Joining tree.

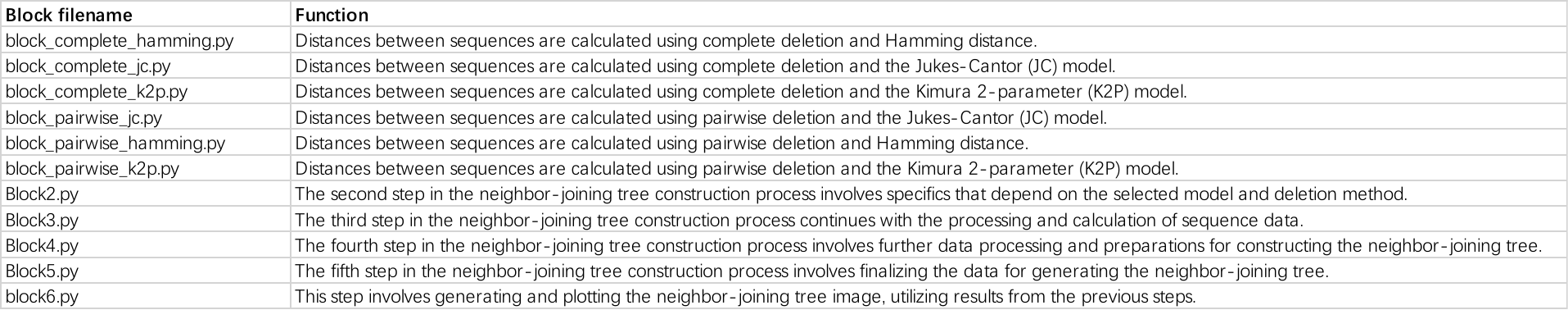

**Supplementary Table2.** Overview of NJGPT's performance in generating phylogenetic tree plots and Newick format tree files for datasets with different sequence lengths and numbers of sequences.

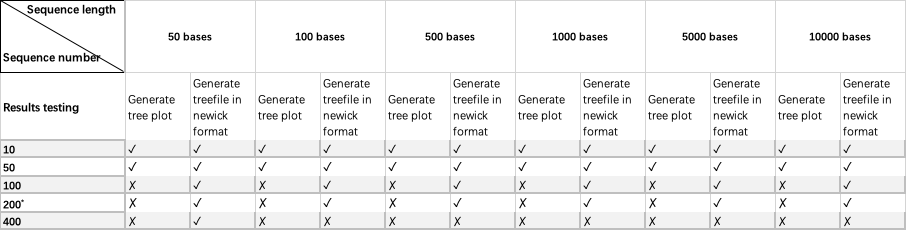

^*^ Indicates that the Newick format tree file generation stutters when the number of sequences is 200, but the user can find the result by refreshing the web page or in the record of NJGPT running block5.

**A**
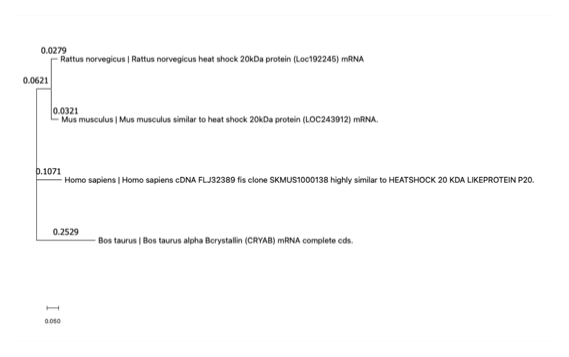

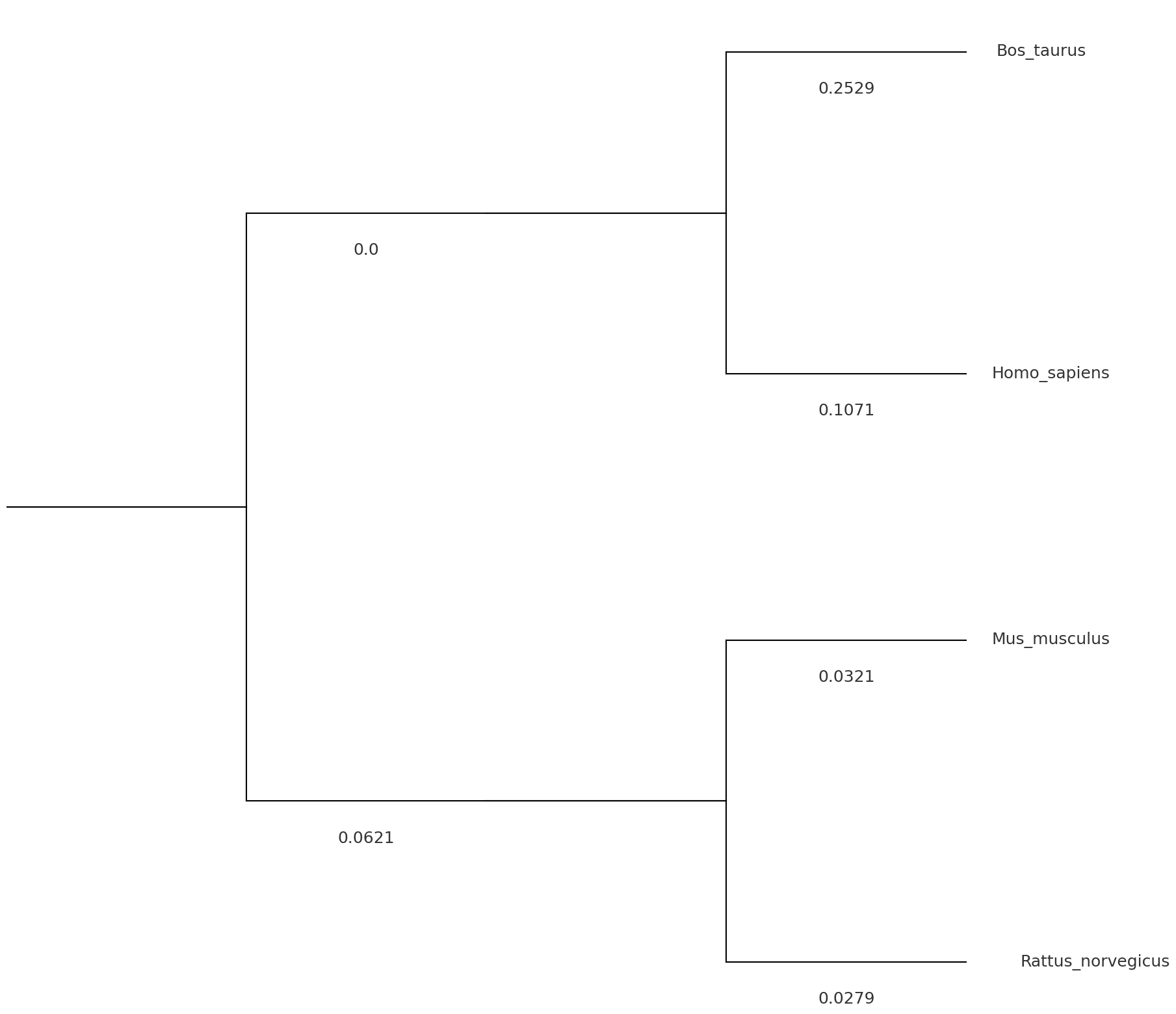

**B**
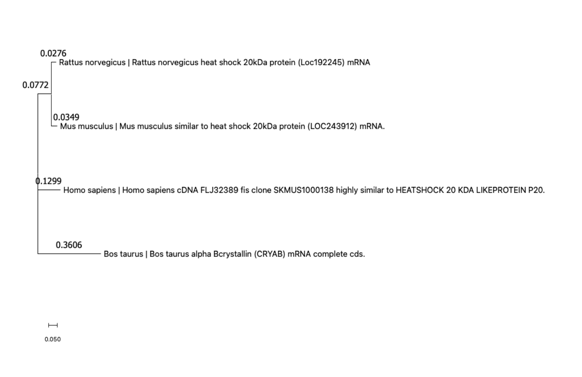

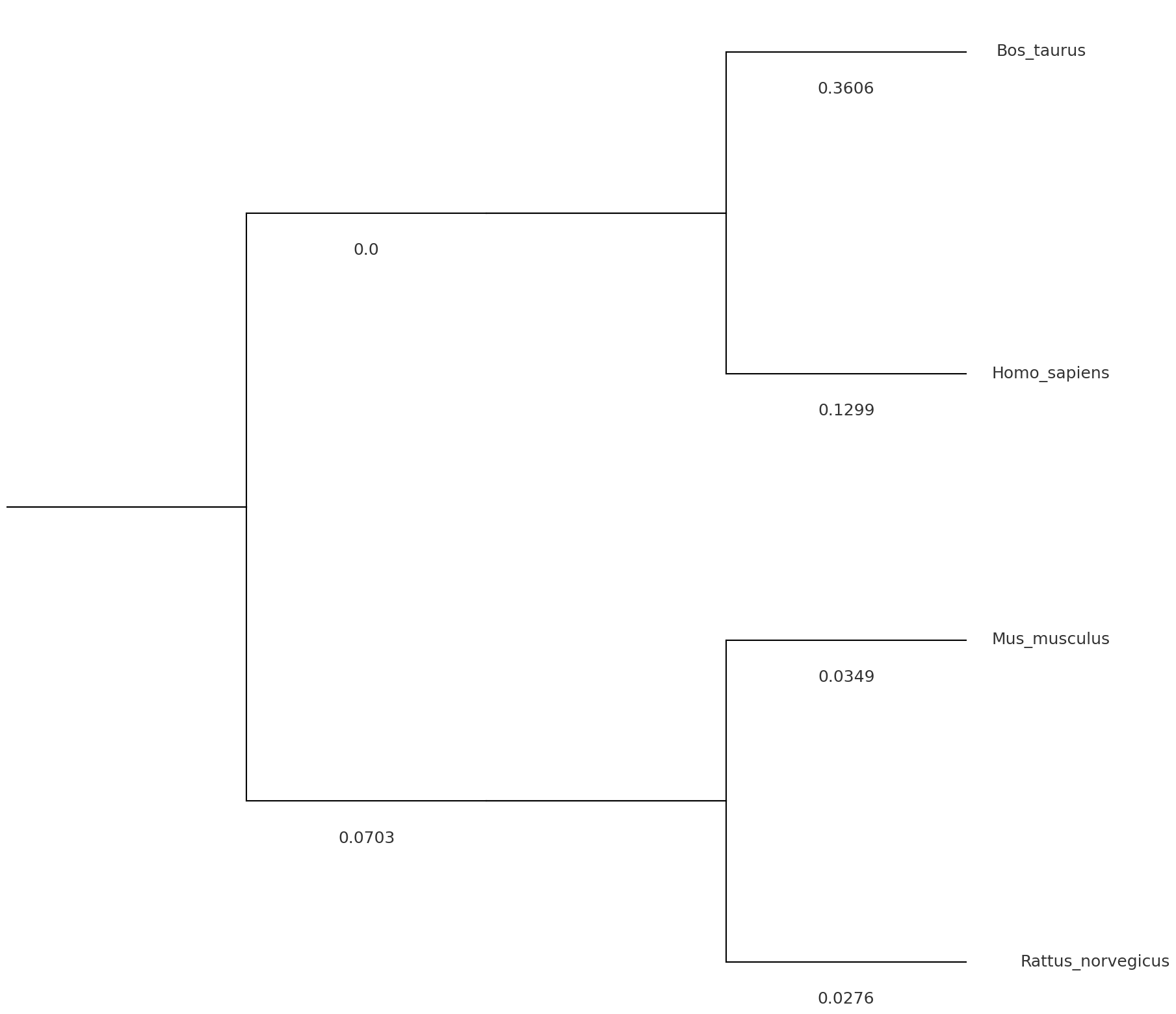

**C**
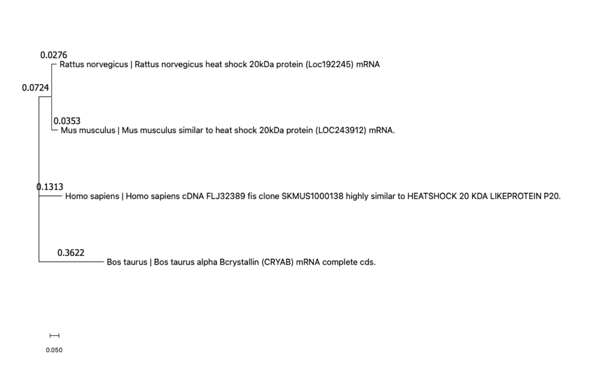

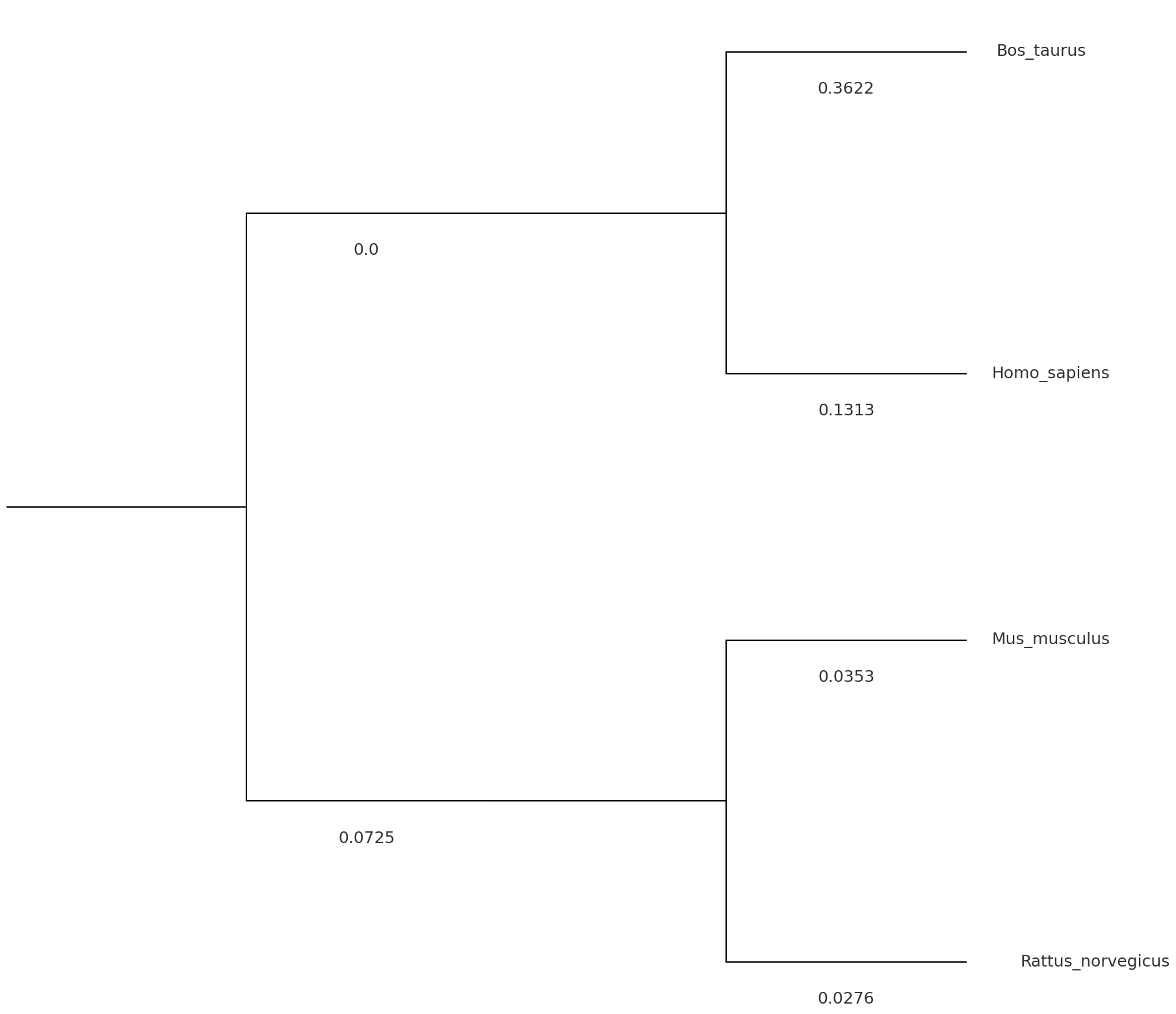

**D**
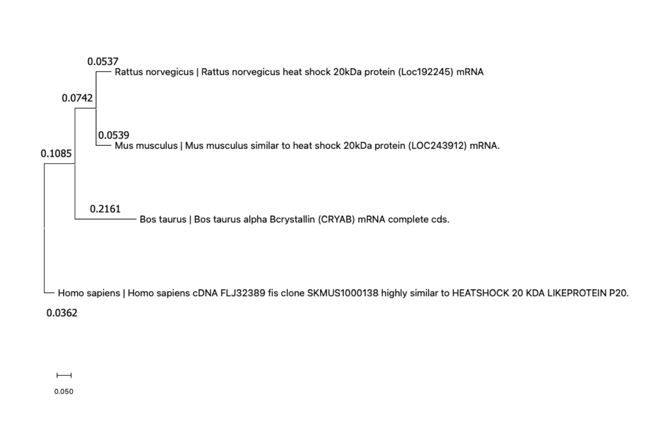

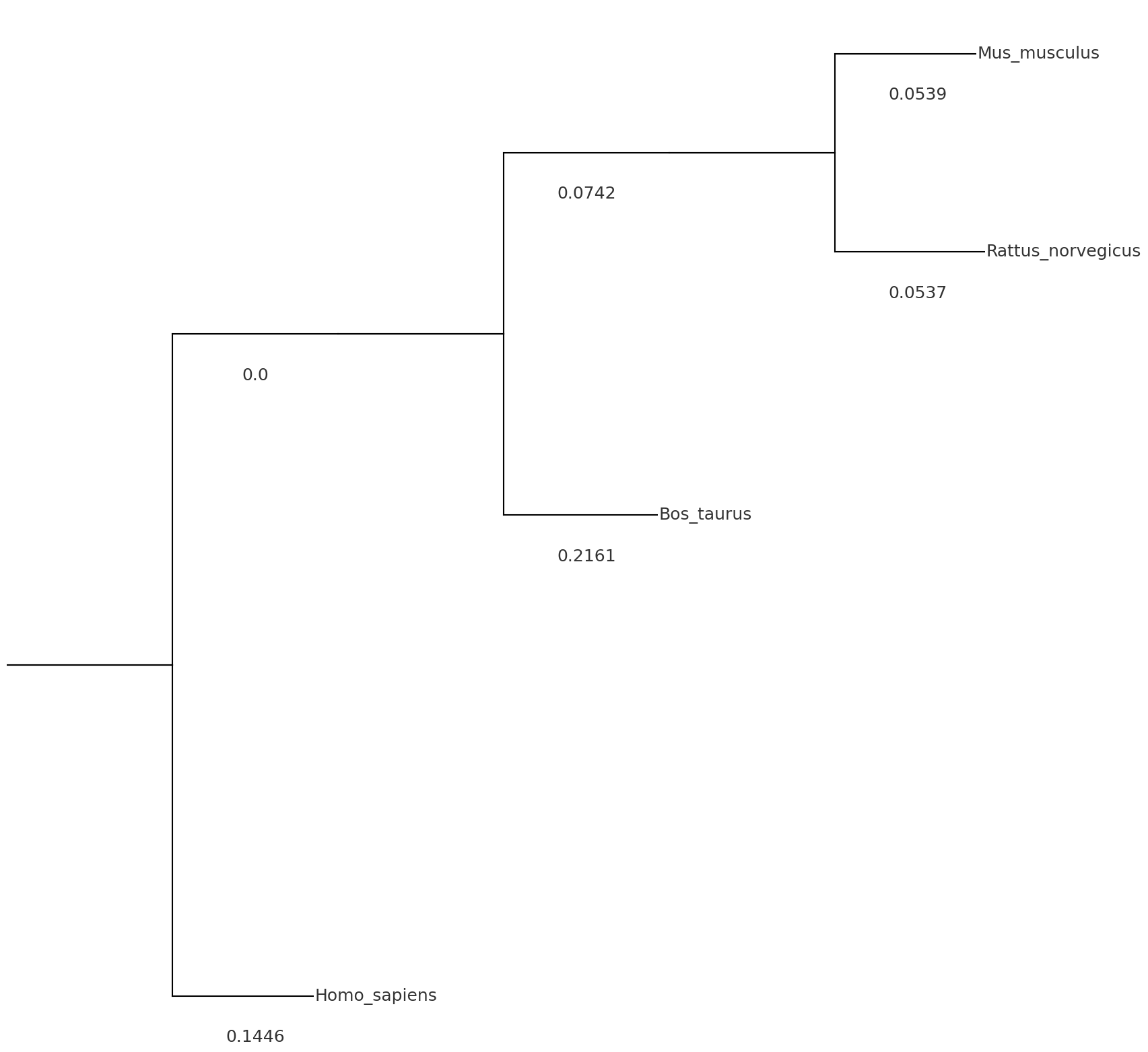

**E**
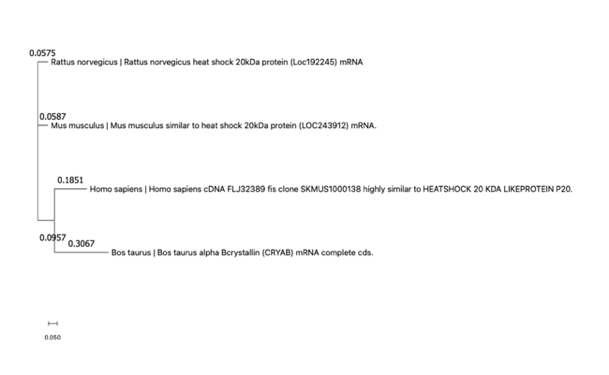

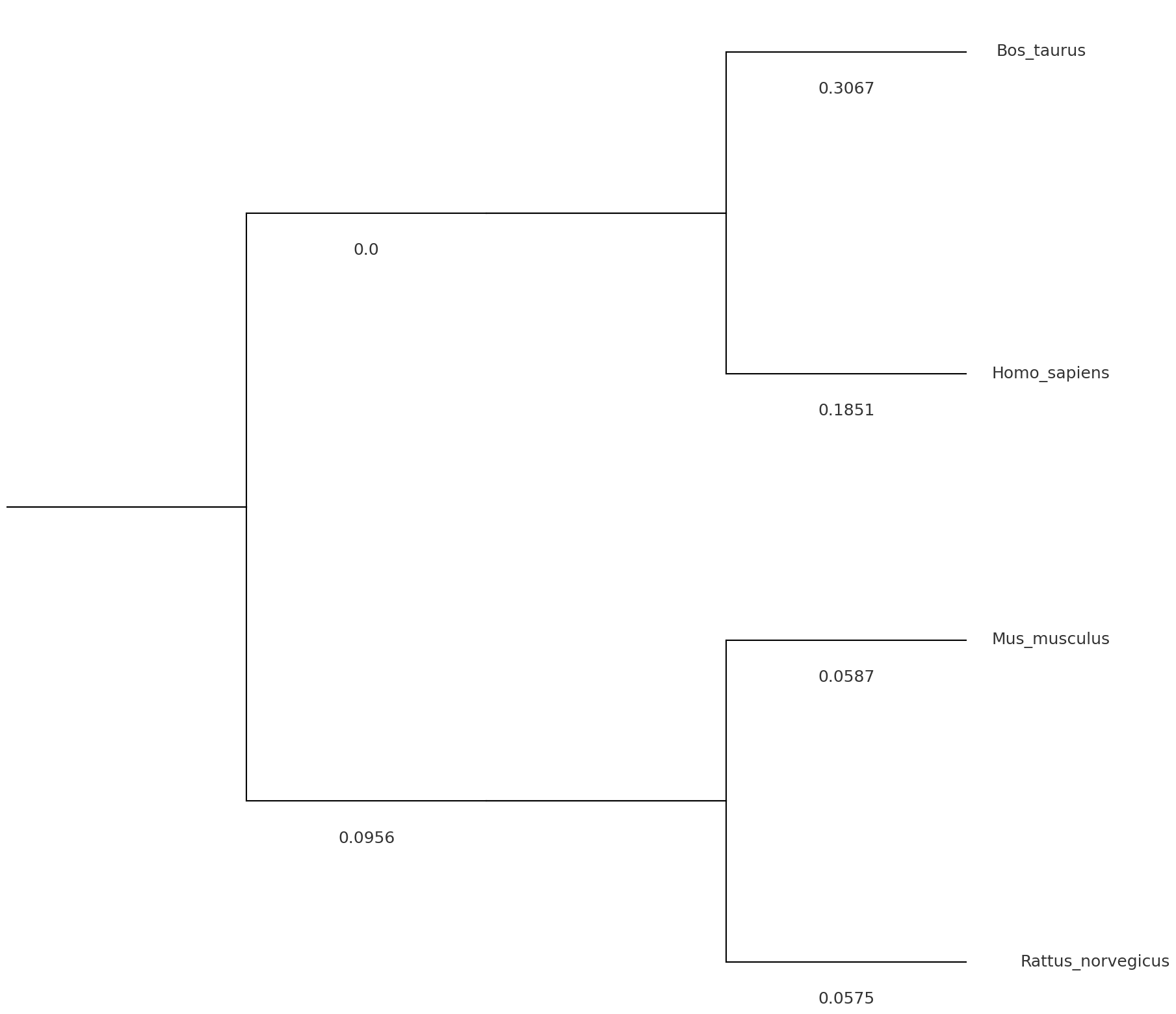

**F**
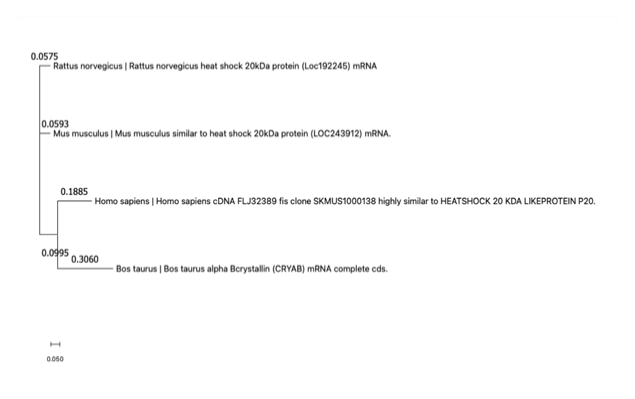

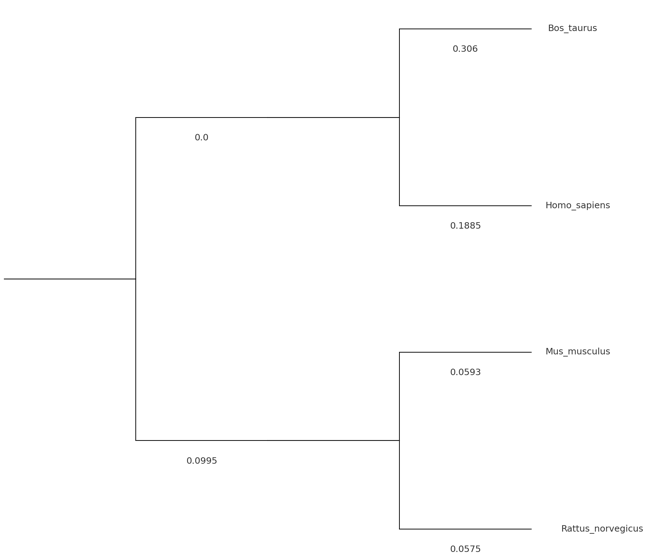

**Supplementary Figure1.** **Comparison of phylogenetic tree reconstruction between MEGA and NJGPT using the “hsp20.fasta” example file across various models.** The left panel displays the results from MEGA, while the right panel shows the results from NJGPT using different methods: (A) Complete deletion with Hamming Distance. (B) Complete deletion with JC model. (C) Complete deletion with K2P model. (D) Pairwise deletion with Hamming Distance. (E) Pairwise deletion with JC model. (F) Pairwise deletion with K2P model.

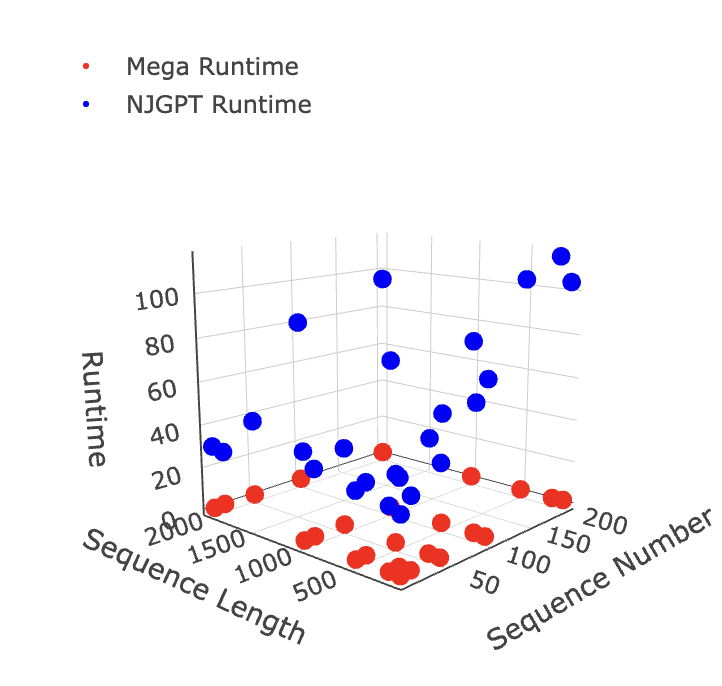

**Supplementary Figure2. 3D plot showing the relationship between run time of MEGA (red) and NJGPT (blue) with the sequences number (x-axis) and sequence length (y-axis).** The run times (seconds) of both MEGA and NJGPT are not greatly affected by increases in sequence length, but NJGPT's run time increases significantly as the number of sequences increases.
